# Can large language models reliably extract human disease genes from full-text scientific literature?

**DOI:** 10.1101/2025.07.27.667022

**Authors:** Danqing Yin, Matthew Ka Siu Leung, Darren Wan Ho Pun, Fiona Haixin Chen, Julie Yujin Kwon, Xinyi Lin, Joshua W. K. Ho

## Abstract

Manual extraction of high-fidelity gene-disease-phenotype information from human genetics literature is a labor-intensive task that requires trained human genetics researchers to read through many primary research papers. This presents a major challenge for maintaining up-to-date human disease genetic databases. Recent exploration into large language models (LLMs) opens new directions in automating this manual process. However, most approaches depend on pre-training, finetuning, or specialized generative artificial intelligence (GenAI) tools, but there is a lack of empirical evidence to show whether commercially-available LLMs can be directly used to reliably extract gene-disease-phenotype for human genetic diseases. Herein, we perform a benchmark of the use of three zero-shot prompted LLMs, namely GPT-4, DeepSeek and Claude, without task-specific fine-tuning, to extract human genetic information directly from full text of scientific papers. Using known congenital heart diseases (CHD) genes found in the open access CHDgene database (https://chdgene.victorchang.edu.au/) as the benchmark data set, GPT-4o achieved overall 88.8% extraction accuracy across 23 gene entries containing over 57 references, with 100% accuracy in gene name, 78.3% and 76.7% in disease and phenotype fields respectively. This work introduces a lightweight, easy-to-deploy, and yet robust LLM-based agent named GeneAgent, analyze sources of disagreement, and highlight the feasibility of integrating powerful LLM into genetic evidence synthesis workflows.

**Highlight:**

- First systematic benchmark of LLMs for extracting human gene–disease–phenotype relationships from full-text biomedical articles
- GeneAgent: a lightweight, highly accurate prompt-only LLM agent
- New domain task-specific evaluation framework

## Introduction

The growing pace of human disease genetic research has resulted in an unprecedented volume of scientific literature, the majority of it containing essential information about gene–disease associations, phenotypes, variants, and underlying pathological mechanisms.

There have been various attempts to benchmark the use of generative AI systems to extract gene–disease–phenotype relationships from full text biomedical literature in human genetics. While previous study (Chen et al., 2025) does not directly investigate the use of LLM in extracting gene-disease-phenotype relationship, it applies relation extraction on datasets e.g. ChemProt and DDI2013, which involves identifying relationships between protein-protein and drug-drug interactions with the use of F1-scores for relation extraction to provide benchmark for comparison with zero-shot, few-shot, fine-tuning performances of GPT-3.5, GPT-4, LLaMA 2, PMC-LLaMA on 12 Biomedical Natural Language Processing (BioNLP) datasets. Another study (Chen et al., 2023) also investigated the use of ChatGPT on a broad set of BioNLP tasks, including relation extraction by evaluating GPT-3.5’s performance in extracting ChemProt, Drug-Drug Interaction (DDI) and GAD information using the Biomedical Language Understanding and Reasoning Benchmark (BLURB) framework. Such a benchmarking framework can be adapted or extended for gene-disease-phenotype relationship extraction. Another pilot study (Jahan et al., 2024) also evaluated multiple LLM in six BioNLP tasks including relation extraction on 26 biomedical datasets including BC5CDR, KD-DTI and DDI with BC5CDR being closely similar to gene-disease-phenotype extraction. In addition, LORE is a framework introduced (Li et al., 2025) for producing knowledge graphs of disease-gene relationships mined from PubMed abstracts with 90% mean average precision. A team further assessed the use of GPT-3.5 and GPT-4 in extracting genetic and genomic data from full-text journal articles have (Poretsky et al., 2025), even though the tested literature is on wheat and barley genetics, the methodology applies to extracting gene-phenotype and disease associations from unstructured scientific literature.

What’s more, an automation framework has been proposed (Chang et al., 2024) for extracting disease-gene associations from literature which uses Hit Ratio to measure how well the framework ranks known disease-gene associations. However, it only focuses on abstracts due to length constraints. Afterall, in the field of human genetics, extraction of human gene–disease–phenotype information depends heavily on expert-driven, manual extraction from the scientific paper, often in unstructured text. This is often a labor-intensive and time-consuming process that does not keep pace with the rate of human disease genetic discovery (Hein et al., 2025; Lee et al., 2020).

Automation of the information extraction process using artificial intelligence (AI) methods is important. Traditional text mining in biomedical research mostly relied on classical natural language processing methods (Rebholz-Schuhmann et al., 2012). These systems employed patterns, dependency parsing, and classifiers to extract genetic associations from abstracts and titles in literature repositories such as PubMed Central. While reliable, interpretable, and causally aware AI agents are yet to be developed to accelerate biomedical knowledge discovery (Gao et al, 2025; Ayesha et al, 2025; Li et al., 2025). Traditional biomedical NLP models are limited by their sentence-level scope, reliance on annotated corpus, and inability to reason over full documents or multi-hop relationships (Zheng et al., 2024; Lamurias et al., 2025). Subsequent advances in BioBERT— a fine-tuned large pretrained language model on biomedical corpora (Lee et al., 2020) has improved particularly on named entity recognition (NER) and relation extraction (RE) and full document processing.

Recent emerging techniques in large language models (LLMs) such as GPT-3.5, GPT-4, and BioGPT have opened new possibilities for zero- and few-shot learning in biomedical information extraction (Gill et al., 2024). These models demonstrate superficial fluency and task generalization, including for entity and relation extraction without task-specific training. Several studies have revealed the potentials of LLMs for tasks such as drug–gene–cancer triad extraction (Lai et al., 2025), gene–trait retrieval from plant literature (Poretsky et al., 2025), structured evidence extraction from full-text papers (Gao et al., 2025; Li et al., 2025) gene function extraction from full-text papers (Kumar et al., 2025) as well as genetic variant prediction from unstructured text (Ayesha et al., 2025). However, most of these approaches rely on retrieval-augmented generation (RAG), model pretraining, task-specific fine-tuning, or external confidence scoring pipelines—all of which involve engineering complexity, computational cost, and risks of hallucination.

Despite the abovementioned technical challenges, human genetics literature itself is in nature highly sophisticated to parse. Gene names often overlap with common words (e.g., “MAP,” “ACE”), entity resolution is confounded by species-specific synonyms, and phenotype associations are frequently described implicitly or across sentences (Poretsky et al., 2025). Consequently, LLMs struggle to resolve gene–phenotype relations without clearly stated contextual metadata (Groza et al., 2024, Neeley et al., 2025, Kim et al., 2024), which is insufficient or unstructured in the source literature (Hein et al., 2025). These factors affect model accuracy and curator agreement. Furthermore, LLMs are prone to hallucination, where fabricated but plausible-sounding gene–phenotype relations are generated without factual support (Kodikara and Verspoor, 2024).

There is a lack of systematic evaluation of the use of generative AI models to extract human disease genes from the full text of scientific papers. One main bottleneck is the lack of high-quality human-curated gene lists as a ground truth. To address this issue, we used a recently published open-access comprehensive database on human congenital heart disease (CHD) genes, called CHDgene (Yang et al., 2022). CHD is a malformation of the heart present at birth with an incidence of 1% of live births. CHDgene is a comprehensive database containing a human-curated list of 189 CHD-causing genes based on human evaluation and curation of 438 publications. In this study, CHDgene serves as a gold-standard resource for benchmarking. In domains like CHD, there has not been any systematic benchmarking on LLM use in the field of human disease genetics. Our expertise in CHD and one of our authors’ involvement in development of CHDgene provides us with a unique opportunity to perform this benchmarking study. This work is the first of its kind in the field of human disease genetics.

We also introduced a domain-specific benchmark framework, namely GeneAgent, to evaluate LLM-based extraction of gene–disease–phenotype associations from biomedical literature. GeneAgent, a robust, domain-constrained LLM-based agent that reliably extracts gene-disease-phenotype associations from biomedical literature using zero-shot prompting alone. Unlike prior approaches, our method requires no fine-tuning, no RAG, and no ontology integration. This framework includes a multi-dimensional evaluation schema for assessing consistency across gene, disease, and phenotype outputs. To better capture near-matches and address variability in biomedical terminology, we incorporated fuzzy and semantic matching into the side-by-side evaluation process for disease and CHD phenotype fields. The framework also includes a systematic protocol for identifying and categorizing disagreement cases—such as false positives, false negatives, and partial matches—enabling error analysis towards a more realistic and interpretable assessment of LLM performance in human genetics.

## Methods and materials

### Dataset construction and retrieval

To assess the ability of LLMs to extract gene-disease-phenotype information for CHD, we leverage the CHDgene repository (https://chdgene.victorchang.edu.au/),a high-quality, manually curated database. CHDgene integrates published findings of gene-level associations with congenital heart defects and serves as our benchmark for genetic evidence synthesis. We restricted our analysis to protein-coding genes listed in CHDgene, each associated with at least one PubMed-indexed primary research article included under *Selected Reference* section in CHDgene database. Consistency level of information extracted by three state-of-the-art, off-the-shelf, commercially available LLMs—GPT-4o, Claude-Opus-4, and DeepSeek-V3—are quantified across multiple dimensions including gene symbol, gene name, disease associated, and CHD phenotypes.

We have randomly sampled 30 CHD genes from the CHDgene’s gene list, and 23 out of the 30 genes have one or more full-length articles as supporting evidence, the rest of the genes were filtered out. For each of the 23 gene, we have compiled custom webscraping scripts in Python and retrieved the following items (Figure 1):

- The gene symbol and full name
- Disease annotations (e.g., “Heterotaxy”, “TOF”)
- Associated CHD phenotypes
- Indexed biomedical literature with PubMed links

**Figure 1.**
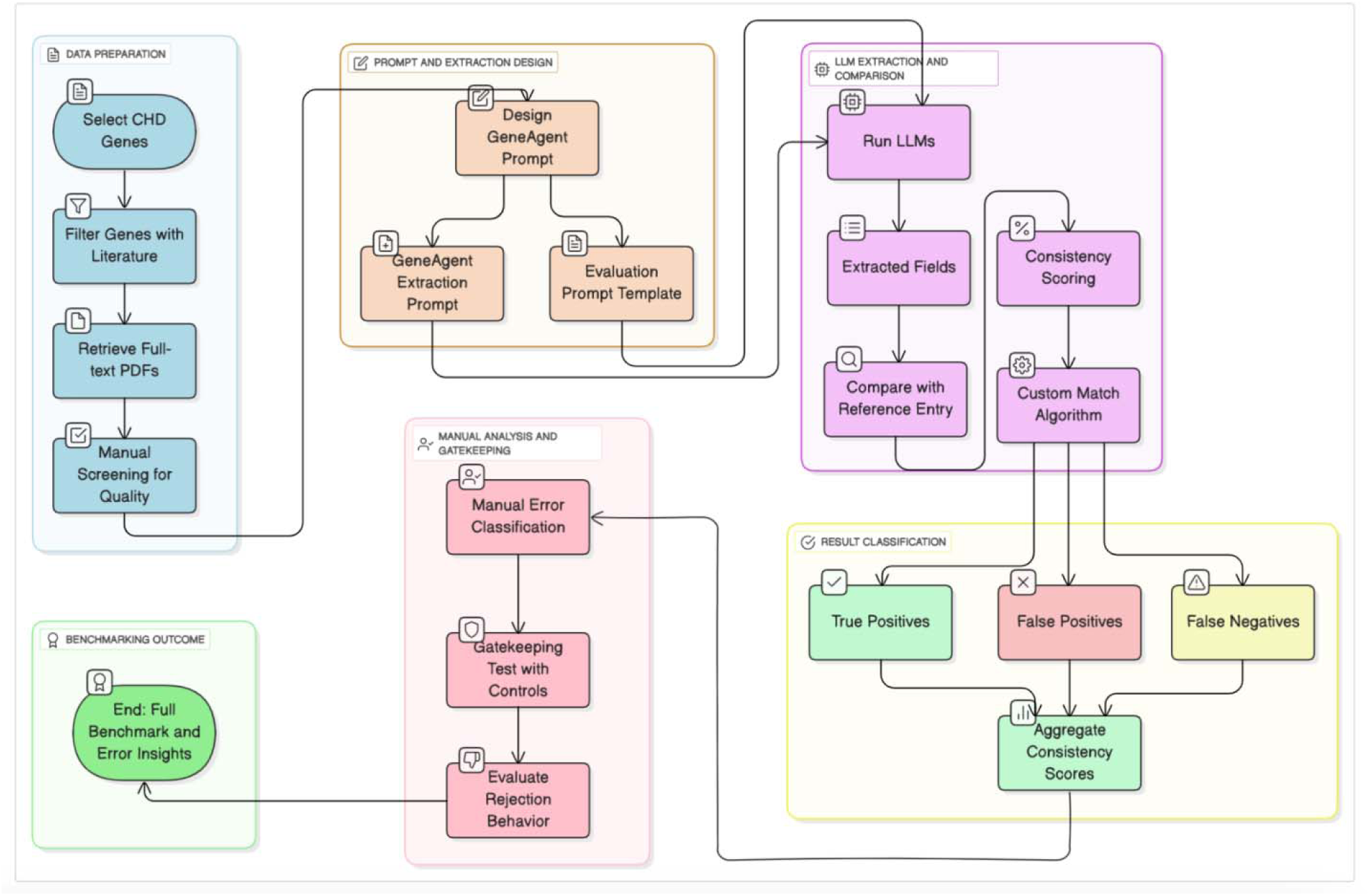
Overall workflow of the GeneAgent study design. Overview of the GeneAgent pipeline for benchmarking LLM-based evidence synthesis of gene– disease–phenotype relationships. The workflow includes data preparation (blue), prompt design (orange), LLM information extraction framework (purple) with result evaluation (yellow), manual error analysis (pink), and final benchmarking (green).

The data are all manually verified for correctness. This benchmark data set contains 23 unique CHD-associated genes, containing a total of 57 scientific papers. This corpus formed the benchmark for evaluating the accuracy of LLM-based extraction.

To construct the test set for LLM evaluation, we retrieved full-text PDFs of the selected literatures from PubMed Central. PDF files were stored and labeled with standardized filenames containing the gene symbol. Manual inspection was performed to ensure PDF quality and relevance by humans. Articles not containing original genetic data or not in English were excluded, ensuring 57 high-quality PDF documents as the final test corpus.

### GeneAgent design

We developed a zero-shot prompt (Figure 2) which is designed to emulate the structure of entries in CHDgene. The goal was to instruct the LLM to extract, in a structured and faithful manner, the gene symbol, full gene name, associated CHD phenotype(s), and disease term (if specified) from the full-text scientific paper.

**Figure 2.**
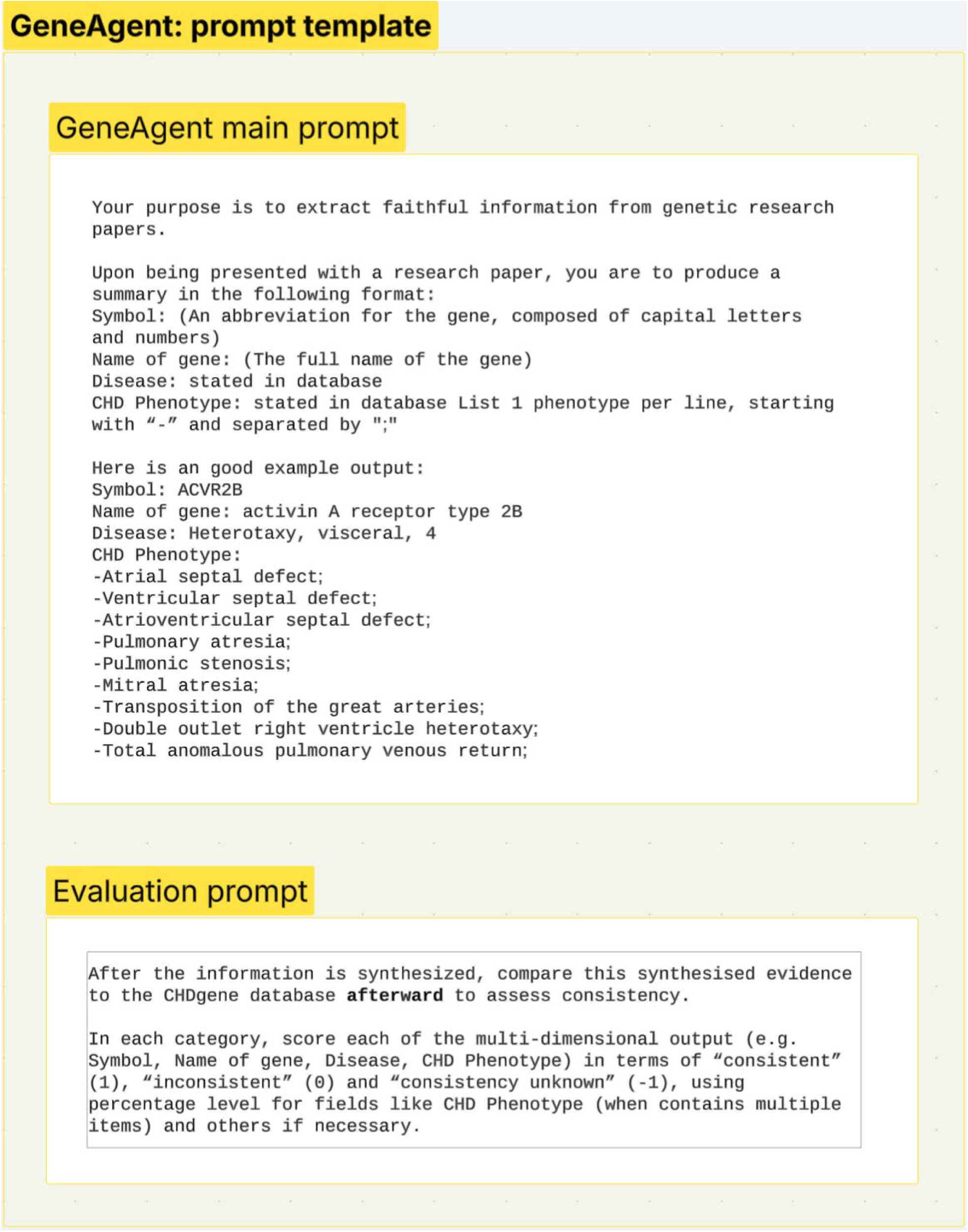
GeneAgent prompt design template.

The prompt consisted of two parts (1) main body of GeneAgent and (2) evaluation. The GeneAgent contains the following components:

- A concise instruction (system message) defining the task of structured evidence synthesis
- An example with a known CHDgene record (e.g., “Symbol: ACVR2B; Name of gene: activin A receptor type 2B; Disease: Heterotaxy, visceral, 4; CHD Phenotype: - Atrial septal defect; - Pulmonary atresia; …”)

Evaluation prompt is as below:

- A set of evaluation guidelines to compare the synthesized evidence extracted from literature against formal CHDgene database records

No retrieval augmentation, external tools, or task-specific fine-tuning were applied.

### Evaluation Metrics

To systematically evaluate the evidence synthesis performance of LLMs, we assessed the structured outputs along multiple dimensions aligned with the CHDgene database schema utilizing the evaluation prompt (Figure 2). For each input paper, the LLM-generated evidence was compared to the gold-standard CHDgene record and scored using a multi-dimensional consistency framework.

### Scoring System: Consistency Evaluation

Each synthesized field — including gene symbol, gene name, disease, and phenotype list — was scored independently using the following criteria:

- Consistent (1): The LLM output exactly matched or was semantically equivalent to the CHDgene entry.
- Inconsistent (0): The output contradicted the CHDgene record (e.g., incorrect gene name, unrelated disease).
- Unknown (-1): The output included plausible but unverifiable information not present in CHDgene or the reference paper (e.g., ambiguous synonyms, inferred phenotypes without direct evidence).

In particular, for CHD Phenotype field, since it contains a list of one or more phenotype terms, was evaluated as follows:

- We computed the percentage of phenotypes correctly extracted compared to the curated CHDgene phenotype list.

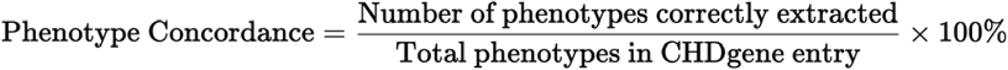
- This score was treated as a continuous metric (0–100%), reflecting partial recovery of phenotype content.

To assess model reliability, we recorded:

- True Positives: Responses containing CHD existing phenotypes supported by the CHDgene entry and the PDF content.
- False Positives: Responses containing CHD phenotypes unsupported by either the CHDgene entry or the PDF content.
- False Negatives: Cases where the reference paper clearly contained CHD phenotype information, but the LLM failed to return any structured output.

Total score equals to the sum of consistency scores across all fields.

### Benchmark design for LLM evaluation

We initially established an exact-match scoring strategy, where perfect alignment of extracted terms with the CHDgene database entries was required for positive scoring. However, recognizing inherent variations in biomedical nomenclature, we progressively enhanced our matching criteria. For the disease dimension, we implemented a fuzzy-matching algorithm using Levenshtein similarity scores (threshold ≥ 85%), accommodating minor differences such as inheritance notations (e.g., autosomal dominant denoted as “(AD)”). .

In the beginning, phenotype terms were standardized by lowercasing, punctuation removal, and singular/plural harmonization (e.g., “defects” unified to “defect”). Subsequently, we implemented synonym mappings to resolve common abbreviations and acronyms (such as “VSD” to “ventricular septal defect”), thereby reducing false negatives stemming from terminological discrepancies. Phenotype consistency was quantified by calculating the proportion of successfully matched phenotype terms, with a threshold of 80% match rate designating a positive consistency count.

### Negative controls

To ensure robustness, we tested LLM behavior on some negative control papers, which consists of 26 randomly sampled articles from PubMed Central what are not related to CHD or genetics. These samples were selected such that the correct response when presented to the system is that the system will indicate that the paper does not contain any CHDgene related information for extraction.

### Data and code availability

All processing scripts were implemented in Python (v3.12). Our entire pipeline, including prompt templates, webscrapping scripts, evaluation notebooks and evaluated results, is made publicly available at https://github.com/hiyin/GeneAgent.

## Result

### Overall accuracy of three LLMs

To assess overall performance, we aggregated consistency scores across all evaluated fields— gene name, disease association, and CHD phenotype—into a single average score per model. As shown in Figure 3, GPT-4o achieved the highest overall consistency, with an average rate of 88.8%. This performance reflects the model’s superior alignment with the CHDgene gold-standard database across both categorical and list-based biomedical fields. Claude-Opus-4 and DeepSeek-V3 displayed comparable overall consistency rates (69.9% and 66.9%, respectively). This convergence suggests their overall extraction fidelity is limited when compared to GPT-4o. These findings underscore the importance of evaluating LLMs not only on isolated information types but also in terms of integrated biomedical comprehension.

**Figure 3.**
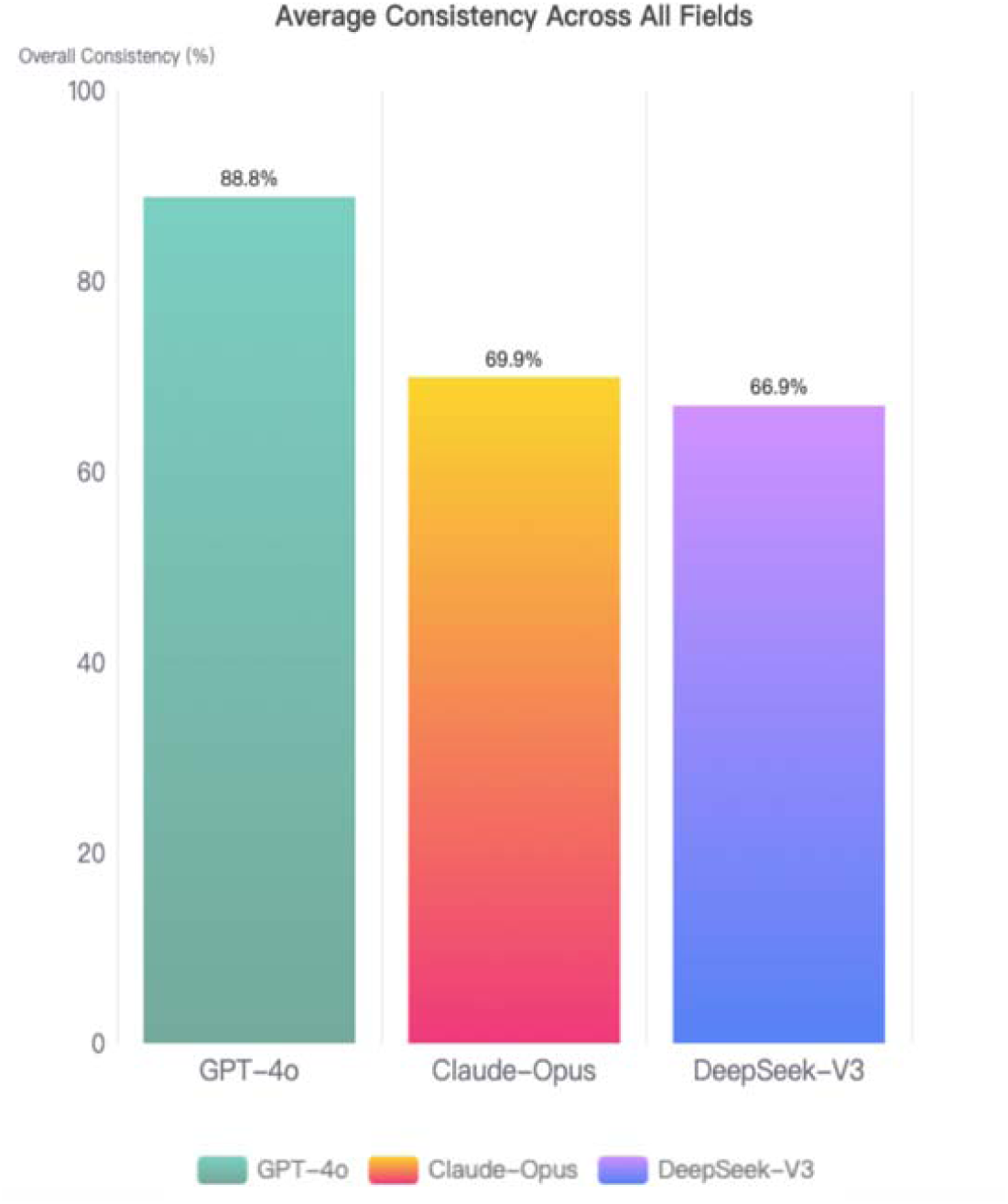
Overall consistency of LLMs in gene–disease–phenotype extraction benchmark. A bar plot showing the average consistency rate (%) of three large language models (LLMs) — GPT-4o, Claude-Opus-4, and DeepSeek-V3 — evaluated across three core categories: gene name, associated disease, and CHD phenotype. Consistency was measured against expert-curated annotations from the CHDgene database. GPT-4o achieves the highest average consistency (88.8%), followed by Claude-Opus-4 (69.9%) and DeepSeek-V3 (66.9%).

### Detailed analysis of LLMs

With respect to multi-dimensional comparison in Figure 4, GPT-4o demonstrated the highest consistency across all dimensions, achieving perfect accuracy in gene name or symbol recognition (100%), and strong performance in disease (78.3%) and CHD phenotype (76.7%) fields. In contrast, Claude-Opus-4 and DeepSeek-V3 both showed moderate performance in CHD phenotype extraction (75.4% and 58%, respectively) but lagged substantially in disease association recognition, with consistency rates of only 30.4% and 28.6%. GPT-4o reached 100% accuracy in name of gene field, followed by Claude-Opus-4 and DeepSeek-V3 scoring 73.9% and 81% closely. Notably, gene symbol extraction appeared relatively robust across all models, whereas disease and phenotype annotations exhibited greater variability.

**Figure 4.**
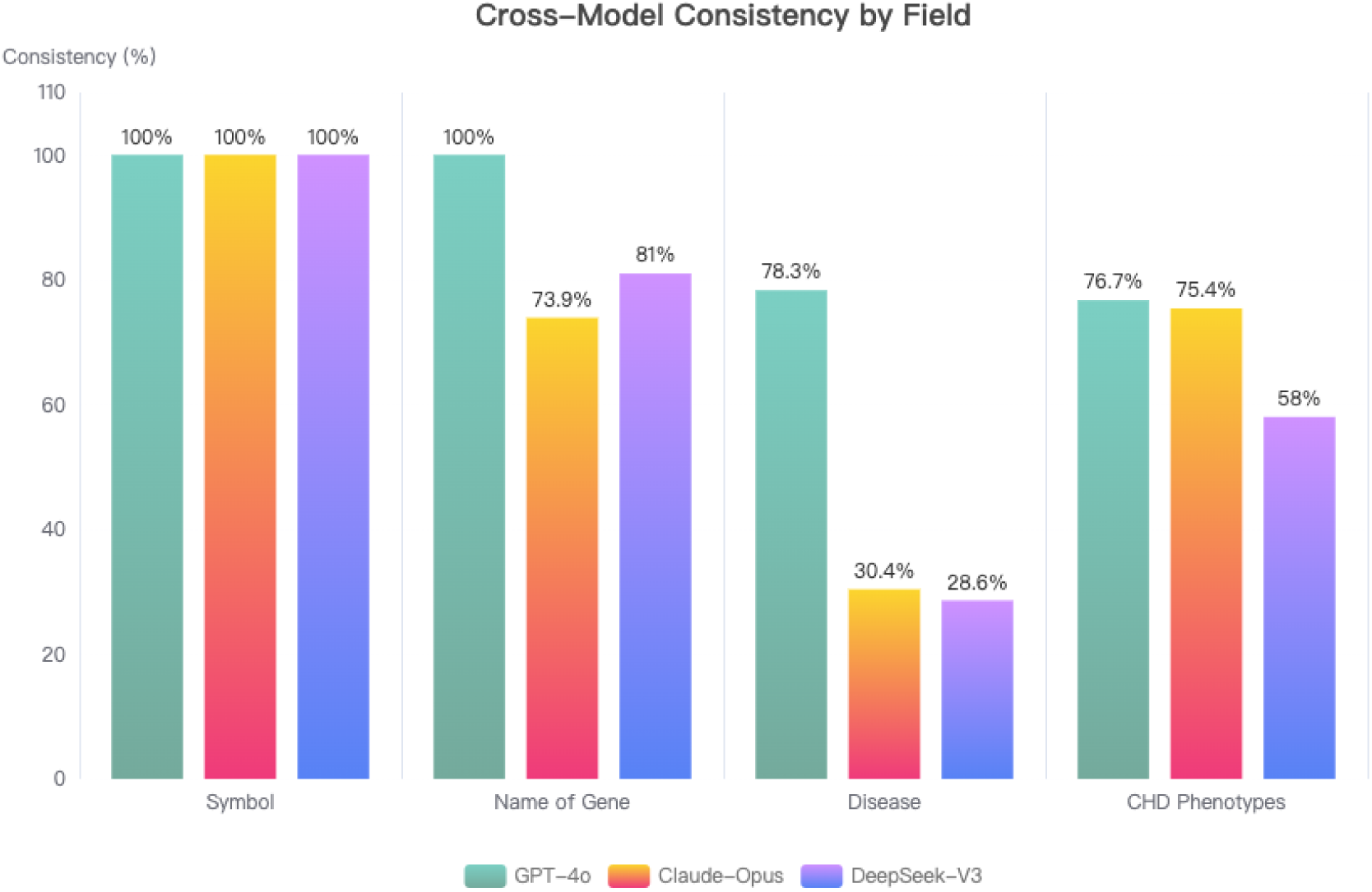
Model-wise consistency of gene–disease–phenotype extraction benchmarked against CHDgene gold-standard annotations. A bar plot depicting the consistency rates (%) of three leading LLMs—GPT-4o, Claude-Opus-4, and DeepSeek-V3. Consistency was assessed across four dimensions: gene names or symbol, name of gene disease associations, and CHD phenotypes, based on comparison against expert-curated CHDgene database entries.

These findings suggest that while LLMs such as GPT-4o are capable of approaching expert-level consistency in certain structured tasks, extracting multi-faceted biomedical associations — especially disease and phenotypes — remains an open challenge for all models. The observed variability highlights the importance of fine-grained benchmarking strategies and domain-specific alignment in biomedical LLM deployment.

### In-depth error analysis: Disease field

One question we ask is which diseases are most or least likely to be wrong under which model, and we detected some patterns, such as length of disease name. Our error analysis has produced both qualitative and quantitative summary on the disease labels below:

It is clear that long, punctuation-rich disease names are far more error-prone (see Table 2). Names that tripped up ≥ 2 models are almost twice as long as universally correct ones (41 vs 20 characters; 5 vs 2 words) and much more likely to contain commas (79 % vs 20 %) or inheritance qualifiers such as “(AD)/(AR)” (86 % vs 40 %). In contrast, short, uncluttered names with minimal punctuation are reliably handled by all models.

**Table 1.** Matching strategies adopted in evaluating the consistency of LLM extracted fields against CHDgene entries.

| Field | Initial Matching | Enhanced Matching |
| --- | --- | --- |
| Symbol | Exact string match | - |
| Name of Gene | Case-insensitive match | - |
| Disease | Lowercase + strip inheritance<br>(e.g. (AD)) | Fuzzy matching (Levenshtein<br>similarity $\geq 0.85$) |
|  |  | Semantic matching |
| **CHD Phenotypes** | Exact match | Fuzzy matching |
|  |  | Semantic normalization |
|  |  | Synonym expansion |

**Table 2.** Lexical features of the 23 benchmarked disease names, stratified by model performance. Median length (characters and words) and frequency of punctuation markers are reported for (i) the full test set, (ii) the 14 diseases that at least two LLMs failed to reproduce, and (iii) the 5 diseases that every model rendered correctly.

| Category □ | All 23 gene-diseases | Diseases missed by ≥ 2 models (n = 14) | Diseases correctly rendered by every model (n = 5) | Significance<br>Welch t test<br>(p-value) |
| --- | --- | --- | --- | --- |
| Median characters | 27 | 41 | 20 | 0.0327*** |
| Median words | 3 | 5 | 2 | 0.0169*** |
| Has commas<br>(%, “ , ”) | 48 % | 79 % | 20 % | N/A<br>(-) |
| Has parentheses<br>(%, “(AD)” /<br>“(AR)”) | 52 % | 86 % | 40 % | N/A<br>(-) |

**Table 3.** Description on textual characteristics of most missed and least missed disease by LLMs.

| Disease | Gene | Description |
| --- | --- | --- |
| **Most-missed (all 3 models wrong) □** |  |  |
| Alagille syndrome 1 (AD);<br>Deafness, congenital ... | JAG1 | 73 chars, 9 words, semicolon<br>& parentheses |
| Leukemia, myeloid/lymphoid<br>or mixed-lineage (AD) | KMT2A | 49 chars, 6 words,<br>parentheses |
| Mental retardation and<br>distinctive facial features (AD) | MED13L | 57 chars, 8 words,<br>parentheses |
| None (ground truth is blank<br>but every model hallucinated<br>a diagnosis) | SPRED2 |  |
| **Least-missed (every model correct)** |  |  |
| KBG syndrome (AD) | ANKRD11 | 16 chars |
| CHARGE syndrome (AD) | CHD7 | 19 chars |
| Costello syndrome (AD) | HRAS | 22 chars |
| Koolen-De Vries syndrome<br>(AD) | KANSL1 | 27 chars |
| Myhre syndrome (AD) | SMAD4 | 25 chars |

Welch two-sample t tests show that disease labels missed by ≥ 2 models are significantly longer than those rendered correctly by all three models, both in character count (p ≈ 0.033) and word count (p ≈ 0.017). There are no significances detected for the other two categories (Has commas and Has parentheses).

We observe a clear monotonic relationship between disease-name length and extraction failure—-longer disease name labels, more errors: adding ∼20–25 characters raises the expected error count by one model (see Table 2; Figure 5). Error dispersion rises sharply once ≥2 models fail. When two or more systems miss the same gene, the corresponding labels are no longer just longer—they are structurally more complex, often embedding commas, semicolons, or inheritance qualifiers such as “(AD)/(AR)”. These composite strings stretch the upper whiskers of the box plot beyond 100 characters. Conversely, short, canonical names remain safe. The only 4 genes that every model handled perfectly—ANKRD11, CHD7, HRAS, and KANSL1—all carry concise, single-clause disease terms. Their ground-truth labels cluster tightly around 20 characters and lack additional punctuation, underscoring that brevity and lexical familiarity protect against hallucination.

**Figure 5.**
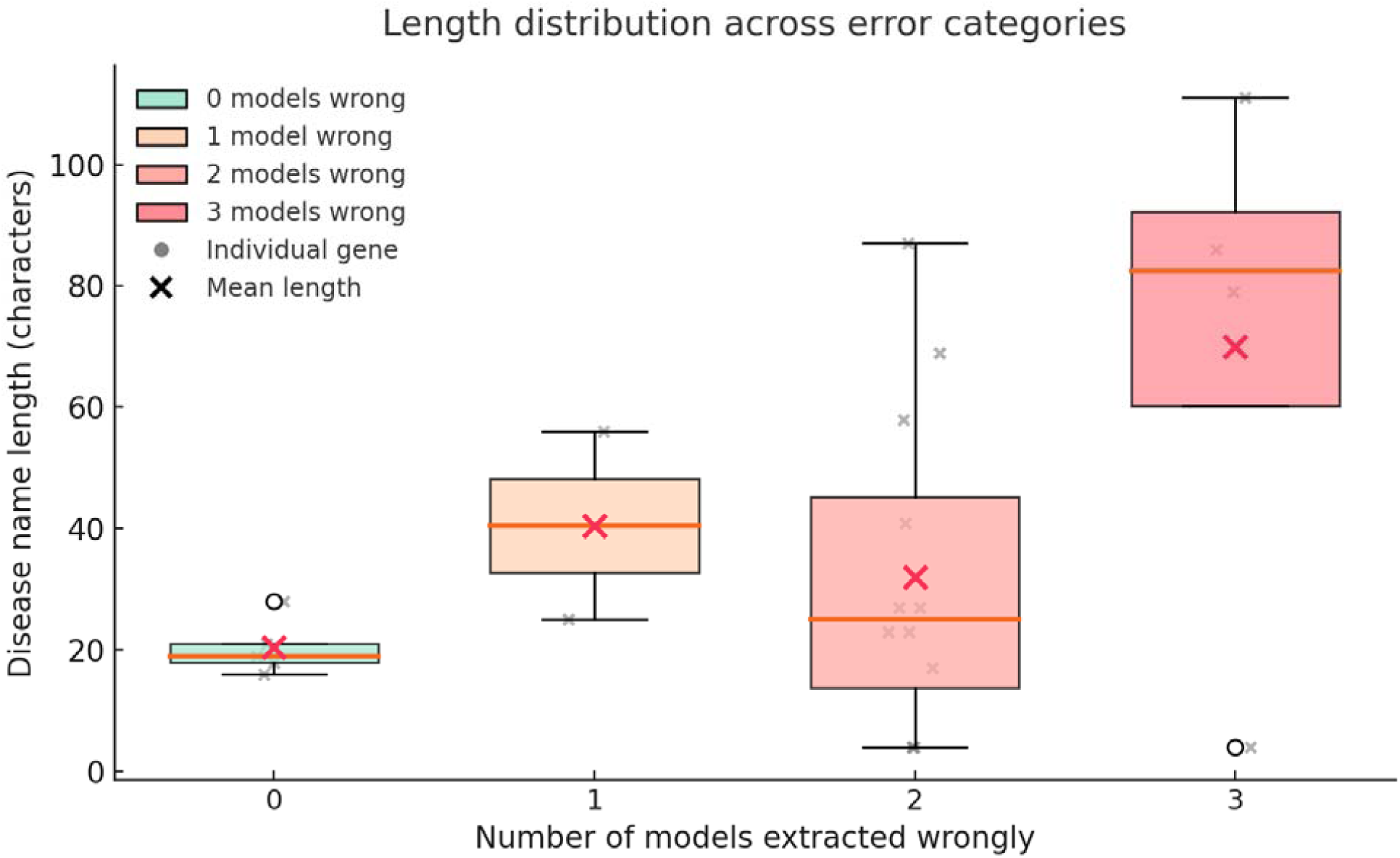
Disease-name length drives extraction errors across models. A box plot that compares the character-length distribution of ground-truth disease names for genes where 0, 1, 2 or 3 large language models (GPT-4o, Claude-Opus-4, DeepSeek-V3) produced a mismatch. Red colours encode the number of models extracted wrongly, grey cross mark each disease name and red “×” symbols mark group means.

### In-depth error analysis: CHD Phenotypes field

Next we ask which phenotypes are most or least likely to be wrong? We counted the CHD phenotypes that are false positives and false negatives. The error analysis (Figure 6) indicates that large language models systematically over-predict highly prevalent septal phenotypes while under-detecting anatomically nuanced or terminologically heterogeneous phenotypes. **Ventricular septal defect** and **patent ductus arteriosus** alone constitute almost 1/3 of all **FP**, implying that when uncertainty arises the models default to “textbook” phenotypes. By contrast, omissions are enriched for complex entities such as heterotaxy, aberrant vascular branches, and subtle valvular malformations, revealing limited recall for less stereotypical descriptors that are likely under-represented in training data. The bidirectional presence of aortic stenosis and patent ductus arteriosus among both false positives and false negatives further underscores inconsistent decision thresholds rather than a unidirectional bias. Collectively, these trends suggest that although there is propensity for LLMs to “hallucinate” these broad commonly cited CHD phenotypes as FP when the source text mentions them tangentially, FN terms are either (i) nuanced variants of common phenotypes (e.g. *mitral stenosis* vs *mitral regurgitation*), or (ii) rarer phenotypes that appear less often in training data.

**Figure 6.**
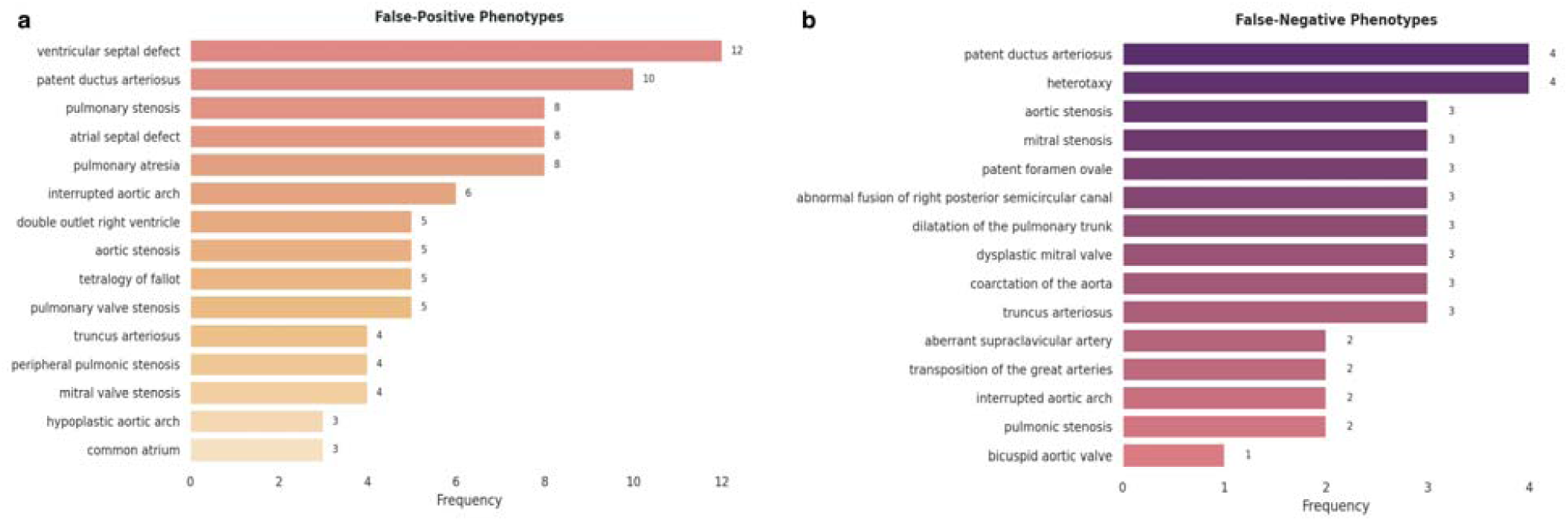
Model-specific distribution of false-positive (FP) and false-negative (FN) CHD phenotypes. Horizontal bar plots show the 15 most frequent false-positive (panel a) and false-negative (panel b) CHD phenotypes produced by the LLM ensemble when compared with the CHDgene reference. Bars are ordered by descending absolute count, with numeric labels indicating the number of errors for each phenotype. Ventricular septal defect (12) and patent ductus arteriosus (10) were the most common false positives, followed by pulmonary stenosis, atrial septal defect, and pulmonary atresia (8 each). False negatives were led by patent ductus arteriosus and heterotaxy (4 each), with aortic stenosis, mitral stenosis, and patent foramen ovale recorded three times apiece. Patent ductus arteriosus and aortic stenosis appeared in both error categories.

**Figure 7.**
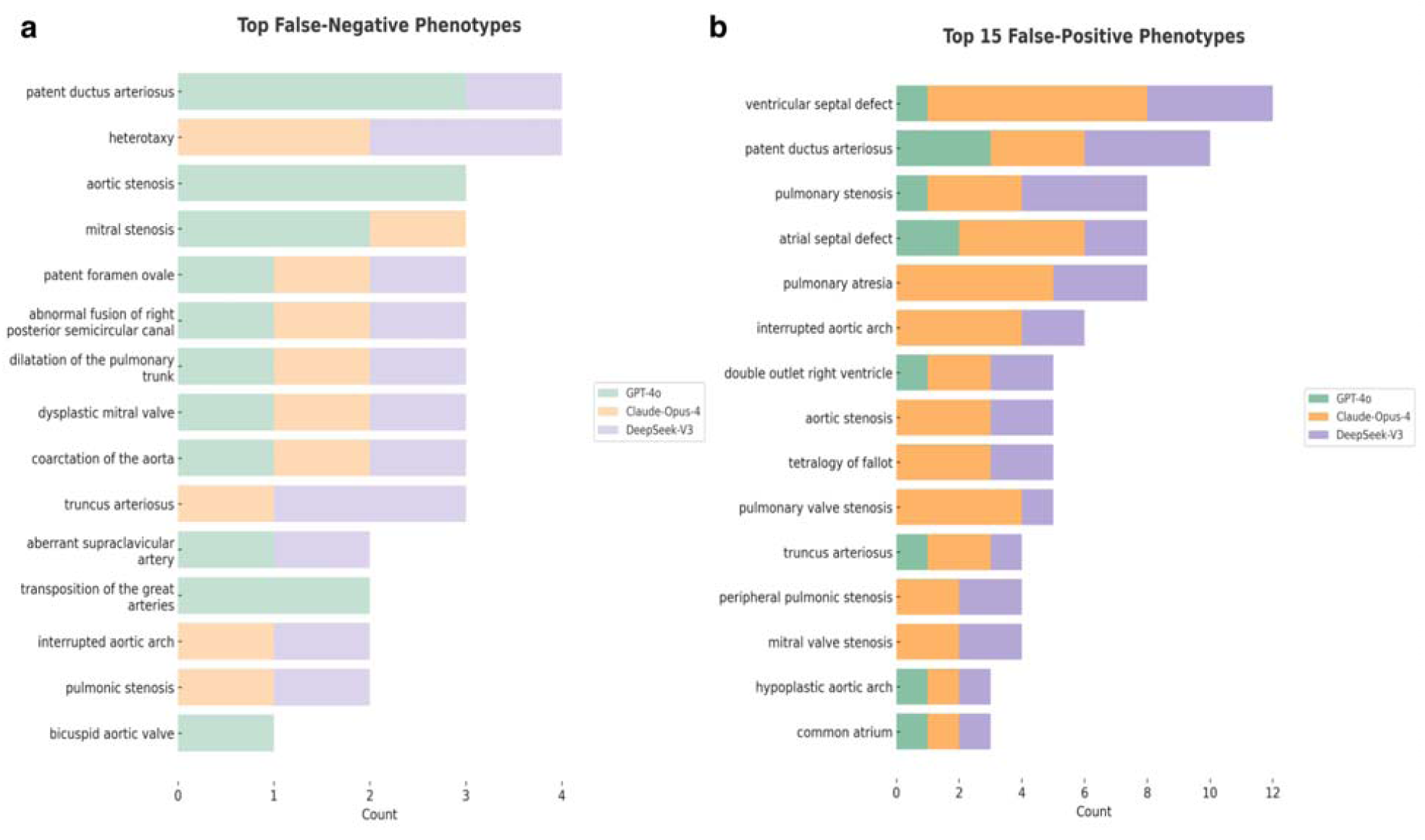
Stacked horizontal bar chart of the 15 most frequent false negative and false-positive CHD phenotypes. Bars are partitioned by model — GPT-4o (green), Claude-Opus-4 (orange) and DeepSeek-V3 (purple) — and ranked by the combined error count for each phenotype. Numeric tick marks on the abscissa denote absolute counts contributed by the three models.

Across the top-15 CHD phenotypes (89 total false positives), Claude-Opus-4 accounted for 48 errors (54 %), DeepSeek-V3 for 31 errors (35 %) and GPT-4o for 10 errors (11 %). The two most over-called phenotypes—ventricular septal defect (12) and patent ductus arteriosus (10)— were driven largely by Claude-Opus-4 (≈67 % and 50 % of their respective totals). A similar pattern was observed for other common septal anomalies (pulmonary/atrial septal defects, pulmonary atresia), whereas model contributions converged for the least frequent items (e.g. hypoplastic aortic arch, common atrium; each three total errors with roughly equal model shares).

### Prompt-specific gate-keeping effectiveness

The zero-shot prompt we used does more than ask the model to list a gene, disease and CHD phenotype; it also lays out a set of gate-keeping rules that must be satisfied before any extraction is attempted. If the provided paper’s PDF file does not contain any human genetic content. Specifically:

- No gene symbols or gene full names are mentioned.
- No references are made to genetic diseases, including CHD.
- There is a complete absence of gene–disease–phenotype relationships or any molecular biology terminology relevant to CHDgene extraction.

As such, the GeneAgent prompt correctly fails this document because it lacks any extractable content within the scope of human genetics or CHD. GeneAgent successfully distinguishes genetics papers from the 26 non-genetics articles by matching against human genetics signatures with 100% gate-keeping effectiveness. True genetics papers repeatedly reference gene symbols (KMT2D, SMAD4), disease, and CHD phenotypes, excluded paper lack of these markers or mentions genes only parenthetically as background.

## Discussion

The emergence of general-purpose LLMs capable of extracting gene–disease–phenotype relationships from unstructured literature presents an inflection point in the curation of human genetics knowledge bases. In this study, we systematically benchmarked three leading commercially available LLMs — GPT-4o, Claude-Opus-4, and DeepSeek-V3 — on a curated corpus of CHD genetics articles, and assessed their capacity to replicate structured outputs across four key fields. Our findings reveal both the immediate potential and the critical limitations of current LLMs in evidence synthesis.

### Fully automatic evidence synthesis of human gene-disease-phenotype relationship remains a challenge

All three LLMs accurately extracted gene symbols and gene names with minimal error. GPT-4o consistently outperformed Claude-Opus-4 and DeepSeek-V3 across all fields. However, accurate disease attribution—particularly fine-grained diagnoses that include inheritance patterns or CHD-specific subtypes — remained challenging for all models, with exact match rates falling below 28.6% (Figure 4) even under relaxed matching criteria.

This trend recapitulates findings from earlier phenotype-focused evaluations (Groza et al., 2024) and underscores the limitations of generalist language models when tasked with clinically granular output.

### Disease and phenotype labels are difficult to extract

The disease field remains the most error-prone output. We identify three dominant causes: (1) Granularity mismatch, where models omit diagnostic qualifiers (e.g., “CHARGE syndrome (AD)” simplified to “CHARGE syndrome”); (2) Lack of ontology anchoring, which prevents synonym normalization (e.g., “Heart and brain malformation syndrome(AR)” vs “SMG9-deficiency syndrome (Heart and brain malformation syndrome, OMIM #616920)”); and (3) Bias toward syndromic generalizations, likely rooted in pretraining corpora that over-represent high-level diagnostic labels (Ji et al., 2023).

The CHD Phenotype field errors exhibit a different but equally systematic pattern. False positives overwhelmingly target “textbook” phenotypes — ventricular septal defect (VSD), patent ductus arteriosus (PDA), and tetralogy of Fallot — regardless of whether these are emphasized in the source paper. This reflects a long-tail prior bias: the over-representation of certain phenotypes in biomedical corpora skews token probability and inflates common terms under uncertainty (Hoffman & Kaplan, 2002; Groza et al., 2024). Conversely, false negatives are concentrated among subtle or anatomically complex malformations — heterotaxy, arch anomalies, valvular defects — often expressed in indirect or diffuse language.

Importantly, many phenotype errors arise not from hallucination in the traditional sense, but from intrinsic co-reference drift. As documented in the clinical summarization literature (Gandhi et al., 2025; Huang et al., 2024), LLMs tend to elevate secondary findings or umbrella terms when a precise causal link is under-specified in text. Our hallucination audit confirmed that >90% of “incorrect” disease and phenotype outputs were present somewhere in the article but assigned incorrect salience by the model. Truly extrinsic inventions—terms absent from the source text— were exceedingly rare, confirming prior taxonomies of hallucination in biomedical natural language generation (Li et al., 2025).

### Divergent model behaviour: decoding mechanisms shapes error profiles

While GPT-4o achieved the most balanced overall performance, model-specific decoding strategies shaped distinct error modes. Claude-Opus-4 contributed over half of all false-positive phenotypes, a consequence of its higher sampling temperature and permissive top-p settings (Anthropic Model Card, 2025), which favour exhaustive recall over precision. By contrast, GPT-4o, tuned with reward models that prioritize calibration and brevity (OpenAI, 2025), exhibited fewer hallucinations but occasionally under-reported less salient phenotypes. DeepSeek-V3 trailed both models in rare disease resolution, particularly in multi-clause or non-canonical expressions, suggesting that training corpus diversity governs model resilience to linguistic variability in human genetics-related terms.

These distinctions matter for future application design. Claude’s aggressive generation may be preferable in workflows where high recall is acceptable, while GPT-4o offers greater accuracy in unsupervised extraction settings.

### Toward reliable model-assisted curation

Our results suggest three practical strategies to bridge the gap between raw LLM output and clinical annotations in human genetics database:

1. **Ontology-constrained decoding**. Anchoring generation to structured vocabularies such as OMIM, Orphanet, or the Human Phenotype Ontology (HPO) improves lexical normalization and reduces semantic drift (Groza et al., 2024; Farquhar et al., 2024).
2. **Probability-based confidence gating**. Biomedical hallucination detectors that leverage output log-probability variance can effectively flag low-certainty candidates for human review (Farquhar et al., 2024), minimizing curation overhead.
3. **Prompt-based role restriction**. Explicit instructions to exclude umbrella terms and extract only the primary CHD diagnosis mitigate over-inclusion and co-reference inflation, as also recommended in recent clinical summarization protocols (Gandhi et al., 2025).

Together, these strategies do not require fine-tuning and can be deployed in a zero-shot manner.

### Limitations and future directions

Several limitations are worth considering. First, our benchmark corpus comprises only English-language papers; extending evaluation to multilingual corpora is essential in a global context as some genetic papers are written in a language outside English. Second, all experiments were performed in a zero-shot setting, improvement in performance can be expected through few-shot, chain of thought prompting, and RAG (Verspoor, 2024). We have not specifically compared our zero-shot setting to any of these settings. Third, our field set was constrained to four core identities (Symbol, Name, Disease, Phenotype); other clinically relevant variables— variant detail, inheritance mode, population-specific effects—remain unexplored. Incorporating these will be critical for end-to-end database curation pipelines.

## Conclusion

The current generation of LLMs demonstrates strong competence in extracting structured genetic knowledge from full-text biomedical literature. Yet they remain fragile in the case of semantic nuance, co-reference dilemma, and clinical specificity. Rather than rely on model scale alone, future advances should prioritize ontology-guided evidence synthesis and context-sensitive prompting. By addressing these painpoints, we are close to building a truly scalable, trustworthy biomedical AI agent for evidence synthesis in human genetics literatures.

## Acknowledgements

We are grateful to the CHDgene project team members for curating the gold-standard database used in this benchmark.

## Author contribution

Conceptualization of the study including the methodology was jointly led by JWK Ho and D Yin. DWH Pun, MKS Leung, F Chen and J Kwon contributed to feasibility study and initial prompt development. D Yin developed and LLM prompts and the benchmarking framework. D Yin performed manual literature screening and completion of LLM output cross-checking. X Lin critically evaluated the results and provided feedback. D Yin and JWK Ho wrote and revised the manuscript.

